# SCHNEL: Scalable clustering of high dimensional single-cell data

**DOI:** 10.1101/2020.03.30.015925

**Authors:** Tamim Abdelaal, Paul de Raadt, Boudewijn P.F. Lelieveldt, Marcel J.T. Reinders, Ahmed Mahfouz

## Abstract

**Motivation:** Single cell data measures multiple cellular markers at the single-cell level for thousands to millions of cells. Identification of distinct cell populations is a key step for further biological understanding, usually performed by clustering this data. Dimensionality reduction based clustering tools are either not scalable to large datasets containing millions of cells, or not fully automated requiring an initial manual estimation of the number of clusters. Graph clustering tools provide automated and reliable clustering for single cell data, but suffer heavily from scalability to large datasets.

**Results:** We developed SCHNEL, a scalable, reliable and automated clustering tool for high-dimensional single-cell data. SCHNEL transforms large high-dimensional data to a hierarchy of datasets containing subsets of data points following the original data manifold. The novel approach of SCHNEL combines this hierarchical representation of the data with graph clustering, making graph clustering scalable to millions of cells. Using seven different cytometry datasets, SCHNEL outperformed three popular clustering tools for cytometry data, and was able to produce meaningful clustering results for datasets of 3.5 and 17.2 million cells within workable timeframes. In addition, we show that SCHNEL is a general clustering tool by applying it to single-cell RNA sequencing data, as well as a popular machine learning benchmark dataset MNIST.

**Availability and Implementation:** Implementation is available on GitHub (https://github.com/paulderaadt/HSNE-clustering)

**Supplementary information:** Supplementary data are available at *Bioinformatics* online.

## 1 Introduction

Cytometry is an established high-throughput technology for measuring cellular proteins at single-cell resolution. In the traditional Flow Cytometry (FC), cells are labeled with fluorescent antibodies that bind specific proteins (Picot *et al.*, 2012). Once excited, these antibodies emit light in correspondence with the targeted protein abundance. These light signals limit the number of potential protein markers as the light spectra will eventually overlap. The advanced Mass Cytometry, cytometry by time of flight, or CyTOF expands the number of markers by using metal isotope antibodies (Bandura *et al.*, 2009). The theoretical upper limit to the number of markers is 100, in practice most CyTOF studies use between 30 and 40 markers (Spitzer and Nolan, 2016). Cytometry, including both FC and CyTOF, has become a vital clinical tool and has been applied to several clinical studies, including, but not limited to: diagnose acute and chronic leukemia (Virgo and Gibbs, 2012), monitoring patients’ immune systems after hematopoietic stem cell transplantations (de Koning *et al.*, 2016), defining biomarkers in case-control studies (Stikvoort *et al.*, 2017), and studying the immune cells differentiation in lung cancer (Hernandez-Martinez *et al.*, 2018).

Cytometry is a high-throughput technology resulting in high-dimensional datasets of millions of cells, representing a major challenge in data analysis. A critical step in analyzing cytometry data is grouping the individual cell measurements into distinct cell populations. Traditionally, FC data was manually analyzed by plotting measured intensities of each pair of markers. This allows researchers to gate distinct cell populations by selecting groups of cells with similar protein expression patterns. Cells are grouped by either positive or negative expression of a marker. However, as the number of markers that can be measured increases, the time required for processing this manual labor tremendously increases. This manual gating process is not even applicable for CyTOF data, with ∼2^40^ gates that need to be analyzed when using 40 markers. Additionally, manual gating is biased by the person performing the gating and suffers from reproducibility issues. It also assumes dichotomic expression of a marker (either negative or positive), and ignores the potential of a marker possessing a gradient pattern.

Consequently, researchers have turned to computational methods for analyzing cytometry data. Clustering is an unsupervised process of grouping data points (cells) by its features (protein markers) into distinct groups (cell populations). Many tools have already been published for the task of clustering cytometry data into cell populations (Aghaeepour *et al.*, 2013; Chester and Maecker, 2015; Weber and Robinson, 2016). These tools can be broadly divided into two categories: dimensionality reduction based, and graph based.

In dimensionality reduction approaches, an algorithm first reduces high dimensional data to fewer dimensions in which the data is then clustered. Reducing to two or three dimensions allows visual representation of high dimensional data, which is otherwise impossible. The archetypical dimensionality reduction technique is Principal Component Analysis (PCA). PCA is limited in its usefulness for cytometry data because it fails to capture non-linear patterns which are characteristic of high dimensional omics data. A popular non-linear dimensionality reduction technique in the single cell community is t-distributed stochastic neighbor embedding (tSNE) (van der Maaten and Hinton, 2008). tSNE analyses local neighborhoods of data points and tries to embed the shape of the high dimensional data onto a lower number of dimensions. Clustering can then be performed on the low dimensional embedding to reduce the computational burden of clustering in high dimensional space. Tools such as ACCENSE (Shekhar *et al.*, 2014), ClusterX (Chen *et al.*, 2016), and DensVM (Becher *et al.*, 2014) are all examples of tools that perform clustering after dimensionality reduction. Non-linear dimensionality reduction methods, like tSNE, suffer from scalability to large datasets. Despite recent improvements of the algorithm, calculating tSNE embeddings becomes prohibitively slow for more than million data points (Maaten, 2014; Pezzotti *et al.*, 2017, 2020). Additionally, tSNE embeddings are stochastic and the resulting global structure of the embeddings for identical data will be different between two runs. This can affect any clustering done in the tSNE dimensions; the stochasticity of the embeddings will make the results less reproducible and less reliable.

Hierarchical Stochastic Neighbor Embedding (HSNE) is a machine learning technique that was introduced to solve the scaling problem associated with tSNE. HSNE transforms large volume of high-dimensional data to a hierarchical set of smaller volumes at representing different scales of the data (Pezzotti *et al.*, 2016; Van Unen *et al.*, 2017). At any scale, the data can be processed, such as making tSNE embeddings to visualize the reduced data and subsequently cluster the data at these scales. HSNE implementations exist in Cytosplore (Höllt *et al.*, 2016) and High Dimensional Inspector (https://github.com/Nicola17/High-Dimensional-Inspector). Cytosplore allows users to cluster the 2D tSNE embeddings of each data scale with Gaussian Mean Shift clustering, remedying the scaling problem as at these scales the volume of the data can be orders of magnitude smaller than the full dataset. Nevertheless, the clustering still suffers from reproducibility and reliability because of the stochastic tSNE step to reduce the dimensionality.

A different dimensionality reduction based tool is FlowSOM, which clusters the data using a self-organized map (SOM) (Van Gassen *et al.*, 2015). Briefly, a SOM consists of a grid of nodes, each representing a point in the high-dimensional space. The grid is trained in such a way that closely connected nodes are highly similar. Each point of the dataset is clustered to the most similar node in the grid. FlowSOM does not suffer from scalability issues, as the computation time is extremely fast (Weber and Robinson, 2016). However, FlowSOM cannot automatically find the correct number of clusters, producing less accurate clustering when cell populations are more similar.

An alternative to clustering in low dimensional space, is to cluster the data in the original high dimensional space using graph based techniques. Graph clustering tools like Louvain clustering in Phenograph (Levine *et al.*, 2015) and X-shift (Samusik *et al.*, 2016) start by finding for each data point the *k* nearest neighbors. The neighborhood graph is then analyzed to find regions with high connectivity, indicating clusters of similar cells. Compared to dimensionality reduction tools, graph clustering tools provide more reproducible, reliable and automated clustering, with a better ability to detect cell populations with relatively few cells. On the other hand, these graph based methods suffers heavily from the scalability to large datasets, exemplified by runtimes for Phenograph and X-shift that exceed 5 hours for a dataset of ∼0.5 million cells (Weber and Robinson, 2016).

Here, we present SCHNEL, a scalable, reliable and automated clustering tool for high-dimensional single-cell data. SCHNEL combines the hierarchy idea of the HSNE transform with a graph based clustering, making graph based clustering scalable to millions of cells. SCHNEL produces fast and accurate clustering of cytometry datasets, as well as different types of high-dimensional datasets such as the popular machine learning benchmark dataset MNIST and single-cell RNA-sequencing data.

## 2 Methods

### 2.1 SCHNEL workflow

We developed SCHNEL, **S**calable **C**lustering of **H**ierarchical stochastic **N**eighbor **E**mbedding hierarchies using **L**ouvain community detection, a novel method for clustering high dimensional data that scales towards millions of cells. It combines the HSNE manifold-preserving data reduction properties with graph clustering to assign each data point to a meaningful cluster, while performing the actual clustering on a reduced subset of the data. It uses the hierarchical information contained in HSNE to assign the predicted cluster labels on a subset of the data, back to the full dataset (Fig. 1).

**Fig. 1.**
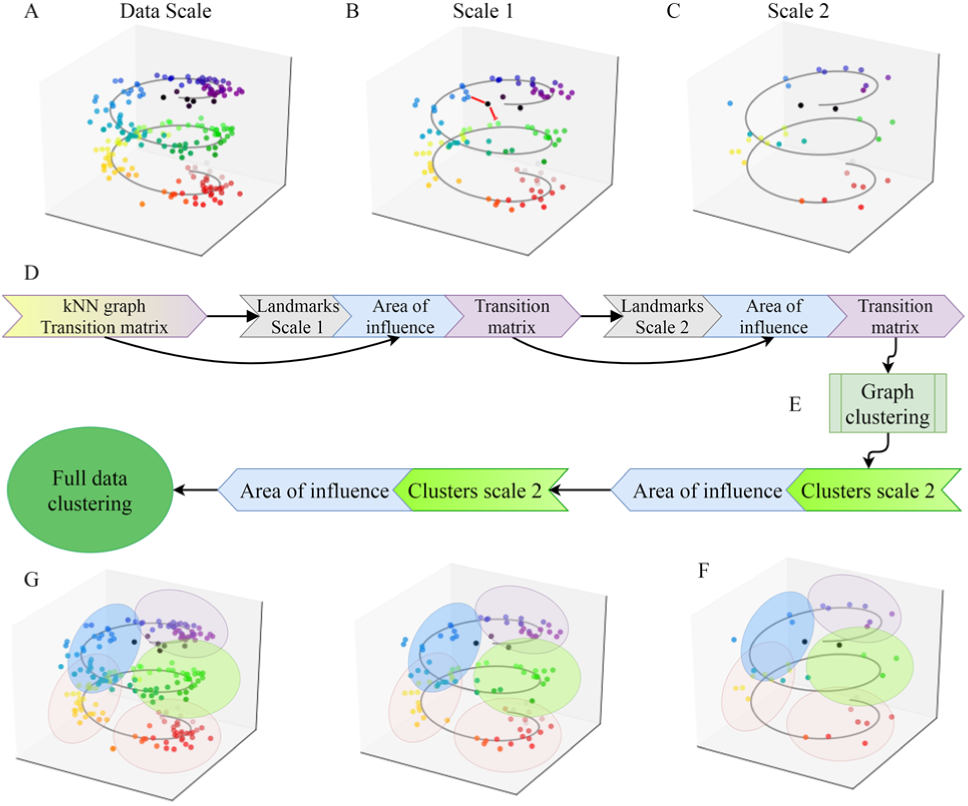
SCHNEL workflow. **(A)** In silico generated dataset: random data points on a spiral manifold in 3 dimensions. The data scale represents all data points within the dataset. The transition matrix is based on the kNN graph of all points. **(B)** At scale 1, highly connected points at the data scale are selected as landmarks. To keep the original manifold, the area of influence for each landmark is calculated, storing the impact/relationship of a landmark at scale 1 on/with the data points at the data scale. The red lines show that performing a kNN at scale 1 will find erroneous neighbors (in Euclidean space), i.e. neighbors that are more distant to each other than when following the spiral manifold. **(C)** Landmarks in scale 2 are subsequently a subset from the landmarks of scale 1. Each scale is sampled from the previous scale in such a way that the non-linear structure of the data in the high dimensional space is retained. **(D)** Flow of information in an HSNE hierarchy. Transition matrices are used to select landmarks for subsequent scales. At all scales (excluding the data scale), the transition matrix is calculated from the area of influence, which in turn is calculated based on the landmark selection process which is derived from the transition matrix at the previous scale. **(E)** Graph clustering is performed on a scale of choice, in this example scale 2. This is a computationally cheap operation since only a small subset of the data is clustered. **(F)** Cluster labels have been assigned to each landmark of scale 2, the labels are now propagated down the hierarchy to the data scale using the information encoded in the area of influence. **(G)** The full dataset now has cluster assignments, while only a fraction of the data was actually clustered.

#### 2.1.1 Creating hierarchy using HSNE

We used HSNE as introduced by (Pezzotti *et al.*, 2016) to construct a hierarchical data representation of the entire high-dimensional dataset. In brief, the hierarchy starts with the raw data, which is then aggregated (summarized) to more abstract scales. At the bottom of the hierarchy, the first scale (data scale) *S*_0_ is the full dataset (Fig. 1A). Using all data points, HSNE begins by constructing a neighborhood graph based on a user defined number of neighbors *k*. Next, HSNE defines a transition matrix *T*_0_ based on two properties. First, the transition probability between two data points, *i* and *j*, is inversely proportional to the Euclidean distance between them. Second, each data point *i* is only allowed to transition to a data point *j*, if *j* belongs to the k-nearest-neighborhood of *i*, otherwise the transition probability is zero. The transition matrix encodes the intrascale similarities between data points.

To define the next scale *S*_1_, HSNE selects representative data points from scale *S*_0_, called landmarks. Landmarks on a given scale *S*_*n*_ represent a subset of (landmark) points at the previous scale *S*_*n*-1_. To find the landmark point at scale *S*_1_, HSNE performs many random walks of fixed length on the transition matrix *T*_0_, starting from each data point at *S*_0_. Next, HSNE records the number of random walks ending at each (landmark) point, reflecting the connectivity of each data point. Data points at *S*_0_ with a connectivity above user defined threshold are selected as landmarks for *S*_1_. As most data points do not meet this threshold, the new scale *S*_1_ is more sparsely populated than the previous scale *S*_0_ (Fig. 1B).

To generate a new scale (say *S*_2_) in the hierarchy, repeating the previous procedure cannot retain the original data structure. For instance, calculating another neighborhood graph on landmarks of scale *S*_1_ will define neighbors with a short Euclidean distance that do not follow the original manifold (Fig. 1B). To preserve the original data structure, HSNE uses a different concept, called the area of influence, to define neighborhoods for landmarks (Fig. 1B-C). The area of influence of a landmark on scale *S*_*n*_ encodes the set of points, from the previous scale *S*_*n*-1_, that can be represented by that landmark. Consequently, the area of influence matrix encodes the interscale similarities, where *A*_*n*_(*i, j*) is the probability that point *i* at scale *S*_*n*-1_ is well represented by landmark *j* at scale *S*_*n*_. The similarities between the landmarks of scale *S*_*n*_ are calculated based on the overlap of their areas of influence on scale *S*_*n*-1_, thus generating the transition matrix *T*_*n*_ for scale *S*_*n*_ (Fig. 1D). As a result, each scale is sampled from the previous scale in such a way that the structure of the full data in the high dimensional space is retained.

#### 2.1.2 Graph clustering

At any scale of the hierarchy, the dataset can be clustered using a graph clustering to define the different clusters of points (cell populations for biological dataset) (Fig. 1E). Depending on the number of (landmark) points on a scale this is feasible or not. In all our experiments all scales, except the data scale, were feasible, as the number of landmark points is at least an order of magnitude smaller than the number of data points in the full data set. We applied the Louvain Community detection, which is a heuristic algorithm that attempts to cluster the graph by maximizing the modularity (Blondel *et al.*, 2008). Modularity is a graph property measuring how well clusters in a graph are separated (Newman, 2006). Clustering is performed on the transition matrices, and the results are propagated to the full dataset using the information encoded in the area of influence (Fig. 1F-G), hence, for all scales, also a clustering of all data points is achieved.

#### 2.1.3 Label propagation

Once the landmarks of a given scale were clustered, these cluster labels are propagated down the hierarchy to label the full dataset. For this task, we used the area of influence (Supplementary Fig. S1). The area of influence *A*_*n*_ at scale *S*_*n*_ is an *i* by *j* matrix, where *j* is the number of landmarks at scale *S*_*n*_, and *i* is the number of landmarks/points at scale *S*_*n*-1_ preceding it. Each row is a probability distribution of point *i* at scale *S*_*n*-1_ being represented by landmarks at scale *S*_*n*_.

We defined a cluster aggregated version of *A*_*n*_ named 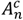, an *i* by *c* matrix, where *c* is the number of clusters obtained from clustering the *j* landmarks at scale *S*_*n*_, and *i* is the number of landmarks/points at scale *S*_*n*-1_. For each row i, the probabilities of landmarks (columns of *A*_*n*_) belonging to the same cluster were summed. The inter-scale connection is defined as the maximum aggregate value of each row in 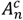. The cluster label each row (data point that needs a label) received was the column (cluster) that had the highest aggregated probability in that row.

#### 2.1.3 Implementation details

We calculated the HSNE hierarchy and converted it to binary format using an adapted version of the High Dimensional Inspector C++ version 1.0.0 that saves the HSNE hierarchy to disk and omits the interactive tSNE. Settings for HSNE were: Beta threshold 1.5, number of neighbors 30, number of walks for landmark selection 200, number of scales round(log10(N/100)) where N is the number of points in the dataset.

The graph clustering is applied using the Python Louvain implementation version 0.6.1 (https://github.com/vtraag/louvain-igraph). The HSNE hierarchy is read using a custom written Python parser. The Louvain algorithm used the transition matrix values as weights, and modularityVertexPartition as maximization objective (Traag et al., 2019).

### 2.2 Datasets

In this study, we applied and evaluated SCHNEL using nine different datasets: one popular machine learning benchmark dataset, seven publicly available cytometry datasets, and one single-cell RNA-sequencing dataset (Table 1).

**Table 1.**
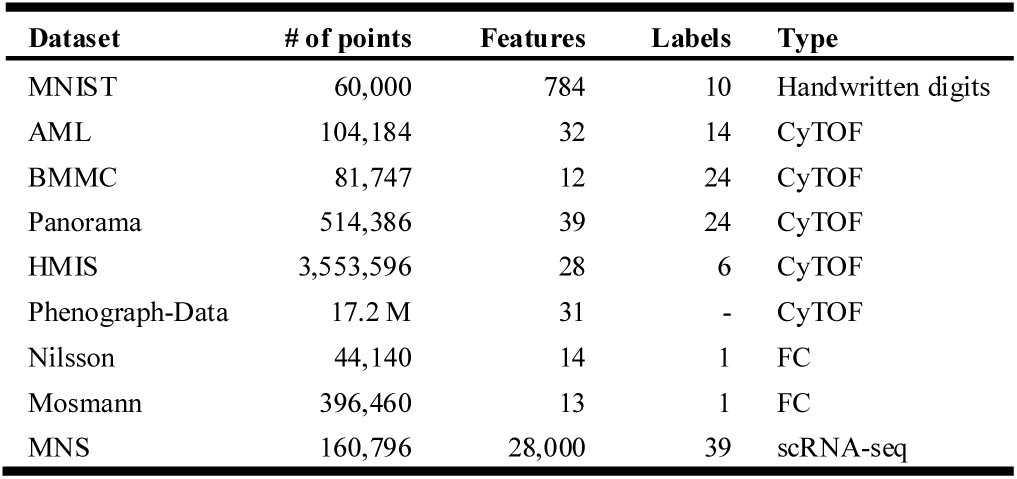
Description of the different datasets employed, showing: the total number of data points (cells or images), the number of features (pixels, proteins markers, or genes, for the MNIST dataset, cytometry dataset and scRNA-seq dataset, respectively), labels indicates the number of ground truth clusters of each dataset, and type of data.

The MNIST dataset contains handwritten digits that were scanned into a computer, each pixel has a value between 0 and 255 and is one of the 784 features of the dataset. It has ten different digits, the numbers 0-9 (http://yann.lecun.com/exdb/mnist/).

All the cytometry datasets are PBMC (Peripheral Blood Mononuclear Cells) or bone marrow samples measured with specific markers to analyze the immune system. The AML dataset is a small benchmark mass cytometry dataset, consisting of bone marrow samples from two healthy humans. The cells were manually gated by experts into 14 different cell populations (Levine *et al.*, 2015). The BMMC dataset is another small CyTOF dataset, containing a healthy human bone marrow sample from a single subject manually gated into 24 cell populations (Levine *et al.*, 2015). The Panorama dataset is a larger CyTOF dataset with 10 replicates of mouse bone marrow samples manually gated into 24 cell populations (Samusik *et al.*, 2016). The HMIS dataset is an even larger CyTOF dataset, consisting of 47 human PBMC samples of healthy, Crohn’s disease, and Celiac’s disease patients. There are no manually gated labels available. The HMIS dataset was analyzed and clustered using Cytosplore resulting into six major immune clusters (van Unen *et al.*, 2016). The largest CyTOF dataset is Phenograph-Data, with more than 17 million cells derived from 21 human bone marrow healthy and leukemia individuals (Levine *et al.*, 2015).

The Nilsson and Mossman datasets are both FC datasets from healthy humans. For both the Nilsson and Mossman datasets there are no full annotations available, only a very small (rare) population is annotated. For the Nilsson dataset, 358 (0.8%) cells are annotated as hematopoietic stem cells (Rundberg Nilsson *et al.*, 2013). For the Mosmann dataset, 109 (0.03%) cells are labelled as CD4 memory T-cells (Mosmann *et al.*, 2014).

Finally, we used the Mouse Nervous System (MNS) single-cell RNA-seq dataset, measuring the transcriptome wide expression of 19 different regions of the MNS clustered into 39 cell populations (Zeisel *et al.*, 2018).

Prior to any analysis, The MNIST dataset and all the cytometry datasets were arcsinh transformed with a cofactor of 5, and all features/markers were used as input to SCHNEL. While the MNS dataset was first log-transformed, next we applied PCA retaining only the top 100 principle components, before inputting the data to SCHNEL.

### 2.3 Evaluation metrics

After propagating the cluster labels to all data points, the clustering can be evaluated using the full dataset. Although the task at hand is unsupervised clustering, we used three different supervised evaluation metrics, as for all datasets, except Phenograph-Data, we had manually annotated cell populations used as ground truth. The evaluation metrics are:

#### The adjusted Rand index (ARI)

measuring the similarity between two sets of cluster label assignments (Rand, 1971). It is adjusted for the chance of coincidentally correctly assigning a pair of data points to the same cluster. It lies in the range of [-1,1], where -1 is worse than random cluster assignment, and 1 is a perfect matching clustering.

#### The homogeneity score (HS)

measuring the pureness of clusters, given a clustering result with a ground truth (Rosenberg and Hirschberg, 2007). It is a score between [0,1], where 1 means that each cluster contains only data points of a single ground truth label.

#### The completeness score (CS)

conversely measuring whether different ground truth groups are captured in distinct clusters (Rosenberg and Hirschberg, 2007). It is also a score between [0,1], where 1 means that all members of a given ground truth label are assigned to the same cluster.

There is a trade-off between high homogeneity and high completeness: e.g. when several ground truth groups all get clustered into one cluster, completeness would be 1 and homogeneity would be 0. It is thus important to evaluate both measures simultaneously.

### 2.4 Benchmarking tools

We benchmarked SCHNEL versus three popular clustering tools for cytometry data, Phenograph (Levine *et al.*, 2015), X-shift (Samusik *et al.*, 2016), and FlowSOM (Van Gassen *et al.*, 2015). Phenograph Version 1.5.2 was used with *k*=30 and default settings for all other parameters. (https://github.com/jacoblevine/PhenoGraph). X-shift was applied using number of neighbors = 20, Euclidean distance, noise threshold = 1, no normalization, no minimal Euclidean length, number of clusters K ranging from 225 to 15. The final number of clusters was determined with the built-in elbow method. Release 26-4-2018 was used (https://github.com/nolanlab/vortex/releases). FlowSOM was applied using x-dim=10, y-dim=10, compensate=False, transform=False, scale=False, maxMeta=40. FlowSOM version 1.1.4.1 was used, available as Bioconductor R package. All experiments were limited to run on a single core Intel Xeon X5670 2.93GHz CPU with 24 GB of memory (to be able to compare runtimes).

## 3 Results

### 3.1 SCHNEL produces meaningful clustering

To evaluate the performance of SCHNEL, we first explored the MNIST dataset, as it has the advantage of easy interpretation of the resulting clusters (recall that the MNIST dataset consists of images of handwritten digits). With SCHNEL, we generated three hierarchical scales of the full data set, and provided a clustering for each scale. Clustering results as well as evaluation metrics are summarized in Table 2. For all scales, SCHNEL produced good clustering, with all evaluation metrics relatively high (> 0.8), despite the difference in the number of landmark points that were clustered at each scale, which vary by orders of magnitude. For instance, scale 3 had only 142 (landmark) data points (∼0.24% of the full dataset) and SCHNEL was still able to produce good clustering with only one cluster less than the original MNIST dataset (9 out of 10).

**Table 2.** Summary of SCHNEL results for MNIST, AML, BMMC and Panorama dataset, across all scales.

| Dataset | Scale | # of points | # of clusters | ARI | HS | CS |
| --- | --- | --- | --- | --- | --- | --- |
| MNIST | 1 | 12,014 | 13 | 0.83 | 0.89 | 0.82 |
|  | 2 | 1,759 | 11 | 0.87 | 0.90 | 0.87 |
|  | 3 | 142 | 9 | 0.82 | 0.85 | 0.90 |
| AML | 1 | 16,031 | 16 | 0.72 | 0.93 | 0.79 |
|  | 2 | 2,595 | 14 | 0.78 | 0.92 | 0.83 |
|  | 3 | 292 | 10 | 0.80 | 0.92 | 0.85 |
|  | 4 | 50 | 6 | 0.94 | 0.85 | 0.98 |
| BMMC | 1 | 13,932 | 14 | 0.92 | 0.88 | 0.94 |
|  | 2 | 2,002 | 14 | 0.90 | 0.87 | 0.93 |
|  | 3 | 118 | 9 | 0.79 | 0.79 | 0.96 |
| Panorama | 1 | 66,466 | 23 | 0.84 | 0.94 | 0.87 |
|  | 2 | 10,943 | 21 | 0.84 | 0.94 | 0.86 |
|  | 3 | 1,436 | 23 | 0.84 | 0.94 | 0.86 |
|  | 4 | 217 | 11 | 0.91 | 0.85 | 0.93 |

Next, we applied SCHNEL to three CyTOF datasets AML, BMMC, and Panorama, and evaluated the clustering of each scale (Table 2). For the AML dataset, SCHNEL provided good clustering of scales *S*_1_ and *S*_2_, with the number of clusters close to the ground truth. While scales *S*_3_ and *S*_4_ showed under-clustering of the AML dataset, probably because there were very few landmark cells at these scales. For the BMMC dataset, SCHNEL under-clustered the data for all scales, with 10 clusters less than the ground truth for both scales *S*_1_ and *S*_2_, but still with relatively good performance. Also, we observed a similar clustering for both scales *S*_1_ and *S*_2_, with an order of magnitude difference in the number of cells between both scales (∼17.04% and 2.45% of the full dataset, for *S*_1_ and *S*_2_, respectively). We obtained similar observations for the Panorama dataset. For scales *S*_1_, *S*_2_ and *S*_3_ (∼12.92%, 2.13% and 0.28% of the full dataset, respectively), SCHNEL produced good clustering, with the number of clusters close to the ground truth. *S*_4_ showed under-clustering as it contained very few landmark cells (0.04% of the full dataset).

The evaluation metrics give a general indication of the clustering quality, but do not show what factors drive the joining or splitting of manually annotated (ground truth) clusters. Therefore, for the interpretable MNIST dataset, we inspected the clustering at the most detailed scale (*S*_1_) and the least detailed scale (*S*_3_) (Supplementary Fig. S2A-B). The contingency matrix at the detailed scale showed good cluster assignments for each digit, although ones and fives were split over multiple clusters (Supplementary Fig. S2A). Further inspection of the average cluster images of the three clusters representing a handwritten ‘one’ (clusters 8, 10 and 11) revealed that their differences relate to the way a ‘one’ is written: straight written ones (Supplementary Fig. S2C), ones written with a slant of 45 degrees clockwise (Supplementary Fig. S2D), and ones written at angles in between (Supplementary Fig. S2E). At the least detailed scale, the split clusters at *S*_1_ were merged, but now also the images representing fours and nines were merged into a single cluster (Supplementary Fig. S2B), due to an overlapping motif between them (Supplementary Fig. S2F).

Next, we checked the contingency matrix of the AML dataset on the most detailed scale *S*_1_ (Supplementary Fig. S3A). The first seven clusters were homogeneous and represent some subsets of the major lineages. In cluster 7, SCHNEL merged CD16 positive and negative NK-cells, and in cluster 9 SCHNEL clumped many of the hematopoietic stem cells (HSPCs). We observed other instances where SCHNEL splits the ground truth classes into multiple clusters. Again, inspecting the cluster averages, which now can be represented as a heatmap of marker expressions, helps to reveal the reasons for splitting or merging ground truth clusters (Supplementary Fig. S3B). For example, clusters 2 and 3 contained almost exclusively CD4 T-cells, which were distinct in their expression of CD7. Clusters 1 and 5 were most different in their expression of CD33. Clusters 4 and 6 were split on distinct expression of CD20. Additionally, cluster 14 contained CD4 and CD8 T-cells with very high expression of CD235ab. Although some of these clusters seem ambiguous, the overall results show the ability of SCHNEL to produce meaningful clusters using only a small fraction of the data (∼15.39%).

For the Panorama dataset, SCHNEL produced almost identical clustering using less cells; at *S*_1_ having 66,466 (12.92%) landmark cells, and at *S*_3_ even with only 1,436 (0.28%) landmark cells (Supplementary Fig. S4). This illustrates the ability of SCHNEL to pick landmark cells that represent the dataset structure well.

### 3.2 SCHNEL outperforms popular cytometry clustering tools

To further evaluated the performance of SCHNEL, we benchmarked SCHNEL against three popular clustering tools for cytometry data (FlowSOM, Phenograph and X-shift), using the three CyTOF datasets: AML, BMMC and Panorama. In terms of the ARI evaluation metric, SCHNEL outperformed other tools across all three datasets, except Phenograph for the BMMC dataset which performed similarly (Table 3). Further, SCHNEL showed better visual partitioning agreement compared to the ground truth manual annotations (Fig. 2).

**Table 3.**
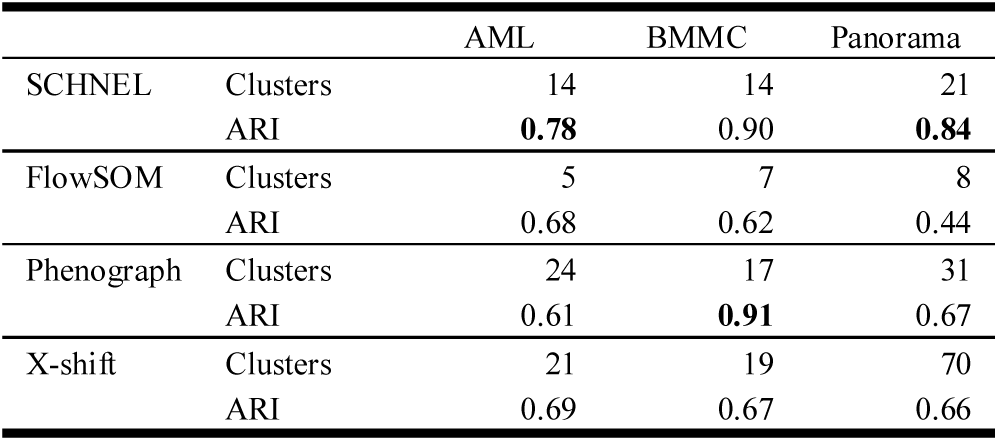
Performance summary of SCHNEL versus FlowSOM, Phenograph and X-shift. Clusters indicates the number of clusters found for each combination of cluster tool and dataset, whereas ARI indicates the Adjusted Rand Index for that combination expressing how much it overlaps with the ground truth data.

|  |  | AML | BMMC | Panorama |
| --- | --- | --- | --- | --- |
| SCHNEL | Clusters | 14 | 14 | 21 |
|  | ARI | **0.78** | 0.90 | **0.84** |
| FlowSOM | Clusters | 5 | 7 | 8 |
|  | ARI | 0.68 | 0.62 | 0.44 |
| Phenograph | Clusters | 24 | 17 | 31 |
|  | ARI | 0.61 | **0.91** | 0.67 |
| X-shift | Clusters | 21 | 19 | 70 |
|  | ARI | 0.69 | 0.67 | 0.66 |

**Fig. 2.**
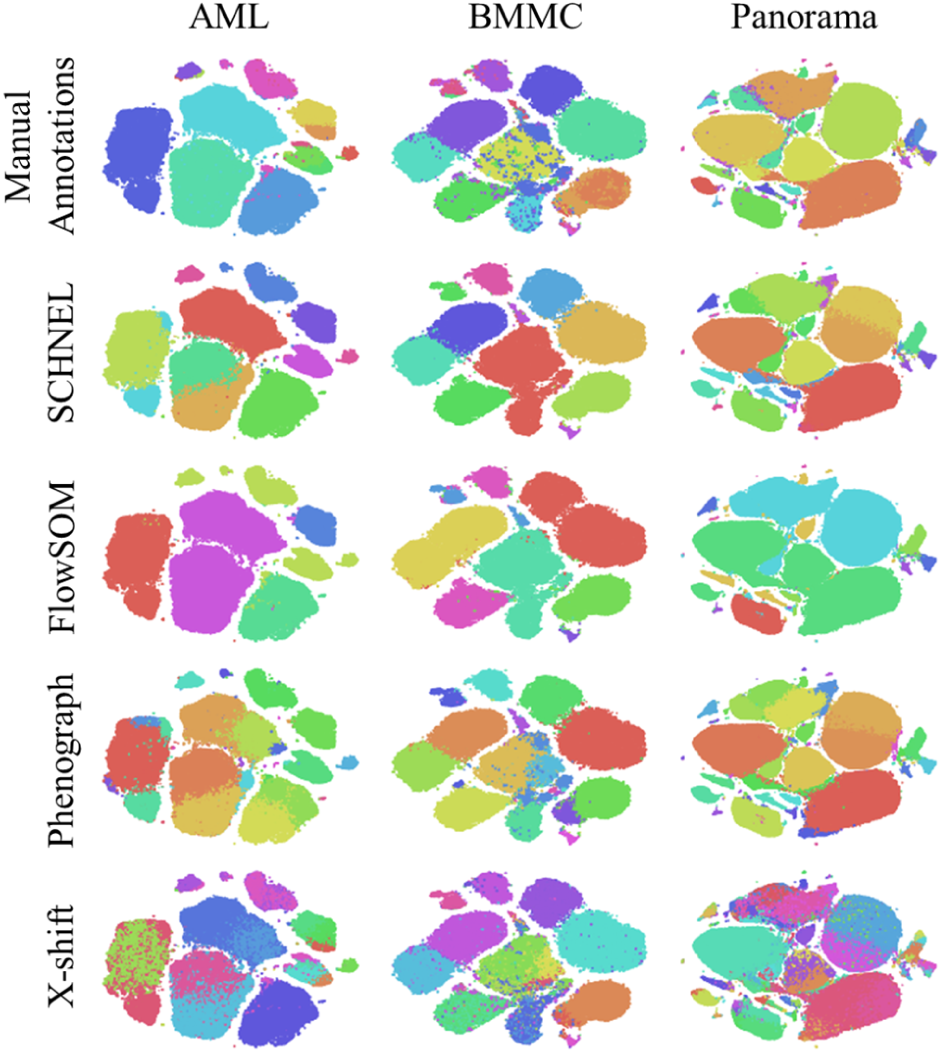
tSNE maps of AML, BMMC and Panorama datasets (columns) colored with different annotations (rows). Manual annotations indicate the ground truth labeling of the datasets. SCHNEL showed good visual agreement with the manual annotations. FlowSOM incorrectly merged different cell populations into mega clusters. Phenograph showed very detailed clustering. X-shift struggled to define clear cluster boundaries.

FlowSOM under-clustered the data using the default settings, in which case FlowSOM determines the optimal number of clusters automatically (Fig. 2). However, FlowSOM was capable of a good clustering when the predefined number of clusters is close to the number of cell populations in the manual annotations (Supplementary Fig. S5). But, generally, this information is not available beforehand. FlowSOM, on the other hand, was extremely fast (clustering the whole dataset under 10 minutes).

Phenograph showed similar partitioning to SCHNEL, but suffered from over-clustering in some cases providing very detailed small clusters (Fig. 2). Speed-wise, SCHNEL was an order of magnitude faster than Phenograph across all cytometry datasets used in this study (Fig. 3). For the 3.5 million HMIS dataset, Phenograph was even not able to complete the clustering after 7 days, at which point it was discontinued.

**Fig. 3.**
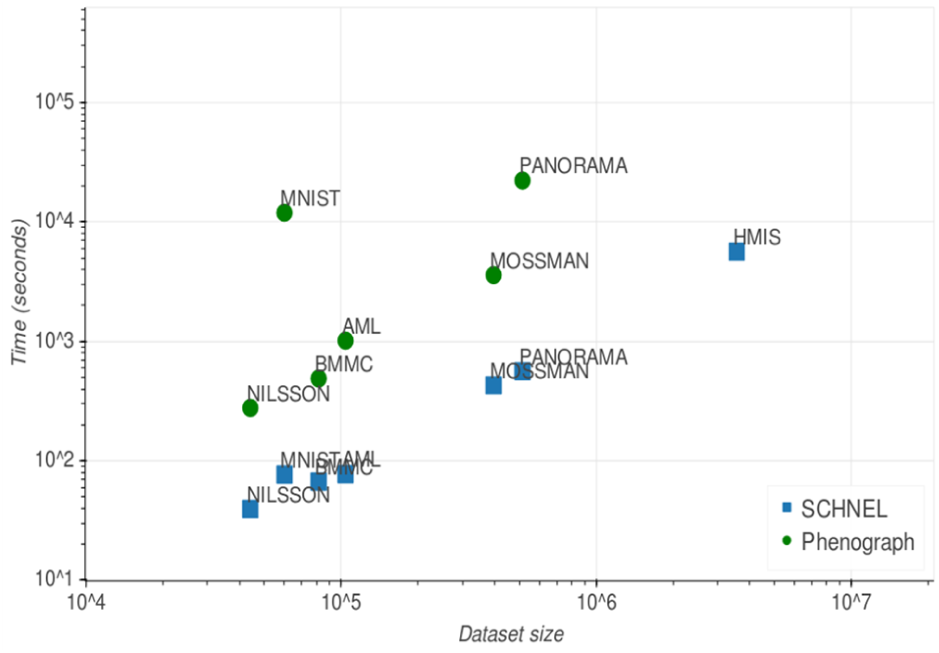
Computation time in seconds of SCHNEL and Phenograph with different dataset sizes. SCHNEL time is the clustering time of all scales in the hierarchy. Axes are log scaled.

X-shift performed reasonably well on the AML and BMMC datasets, but found too many small clusters on the Panorama dataset. Generally, X-shift failed to define clear boundaries between clusters (Fig. 2). X-shift was not timed because its implementation is a graphical user interface, but its computation time was around 30 minutes for the smaller datasets AML and BMMC, and up to 6 hours for the Panorama data. Similar to Phenograph, X-shift was not able to complete the clustering of the HMIS dataset after 7 days.

### 3.3 SCHNEL scales to large datasets

SCHNEL was tested on datasets of different sizes to see how well it scales. Fig. 4A shows the computation time of SCHNEL specified per task. Clustering the most detailed scale (*S*_1_) was the most time-consuming operation. For the HMIS dataset this meant clustering 495,811 landmarks (similar as the entire Panorama dataset). Excluding this scale, the HMIS dataset could be clustered in roughly 50 minutes.

**Fig. 4.**
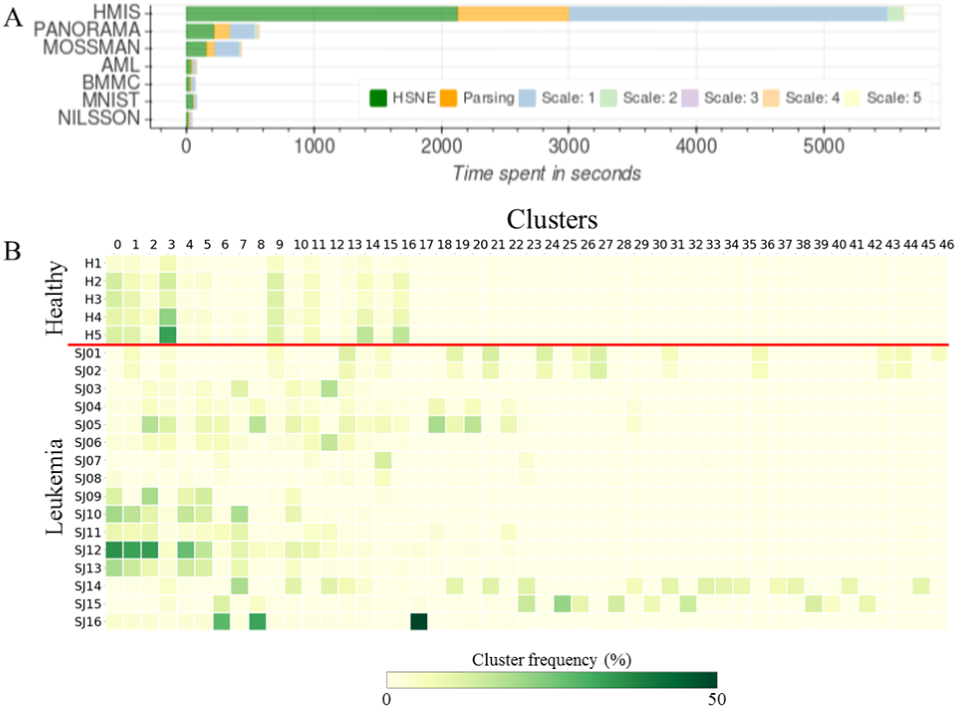
**(A)** Computation time of SCHNEL in seconds for different datasets. Different colors specify different steps in the SCHNEL algorithm. Green is calculating the HSNE hierarchies, orange is reading the HSNE hierarchy into Python, the other colors are times for clustering individual scales. Note that clustering scale 1 of the HMIS dataset (495,811 landmark cells) takes quite some time, showing the benefit of creating hierarchies and (only) clustering at higher scales having less landmarks. **(B)** Cluster frequencies across all the 21 individuals of the Phenograph-Data dataset. Clusters obtained from SCHNEL using scale 5. Red line separates between healthy and leukemia individuals.

Further, we tested the scalability of SCHNEL to cluster the Phenograph-Data with 17.2 million cells, divided over 5 healthy and 16 leukemia individuals. In the original study, this dataset was analyzed per individual using the Phenograph clustering algorithm (Levine et al., 2015). Using SCHNEL, we were able to pool all the cells from all individuals together and obtained a single clustering. We chose to represent the data at six different scales on top of the data scale. These scales contained 2.3M, 378K, 53K, 9K, 784 and 48 landmark cells, from the most detailed scale (*S*_1_) to the least detailed scale (*S*_6_), respectively. We skipped clustering *S*_1_ as it is computationally very expensive. Clustering *S*_2_ to *S*_6_ resulted in 131, 133, 114, 47 and 5 clusters, respectively. Using the 47 cell clusters of *S*_5_, we calculated the cluster frequencies across the 21 individuals (Fig. 4B). Similar to the original study (Levine *et al.*, 2015), we observed a homogeneous pattern across all healthy individuals, while the leukemia individuals had heterogeneous patterns. These results show the scalability of SCHNEL to cluster such large datasets and produce meaningful clustering.

### 3.4 SCHNEL detects rare cell population

Different cell types are expected to have very different abundances and good clustering algorithms should be able to detect rare cell populations which are often interesting to study. We used the Mosmann and Nilsson datasets to test SCHNEL’s sensitivity for detecting small populations. The Mosmann and Nilsson datasets both had manual annotations for only one rare cell population present within their full dataset. The Mosmann dataset contained a population of 109 memory CD4 T-cells. The Nilsson set contained 358 stem cells. Table 4 shows the sensitivity of SCHNEL for detecting these small populations. For both datasets, the cells belonging to the rare populations nicely clustered together at the various scales (CS), but for some scales these clusters also contained many other cells (Cluster size). For the Mosmann dataset, SCHNEL was able to capture the rare population in a single cluster without having many other cells at scales *S*_1_ and *S*_3_.

**Table 4.** Performance of SCHNEL for capturing rare populations in the Mosmann and Nilsson datasets. Cluster size indicates the size of the cluster in which most cells of the rare population were contained. Completeness Score (CS) indicates how many of the annotated rare cells in the original dataset were in the cluster containing most of the designated rare cells.

| Dataset | Scale | # of cells | # of clusters | Cluster size | CS |
| --- | --- | --- | --- | --- | --- |
| Mosmann | 1 | 77,787 | 23 | 181 | 0.94 |
|  | 2 | 9,398 | 18 | 4,949 | 0.99 |
|  | 3 | 1,090 | 14 | 173 | 0.93 |
|  | 4 | 191 | 6 | 110,097 | 0.99 |
| Nilsson | 1 | 9,386 | 23 | 4,314 | 0.93 |
|  | 2 | 1,354 | 17 | 3,269 | 0.93 |
|  | 3 | 125 | 7 | 7,779 | 1 |

### 3.5 Clustering scRNA-seq data using SCHNEL

After showing the potential of SCHNEL to cluster cytometry data, we tested the ability of SCHNEL to cluster scRNA-seq data which has many more features. We applied SCHNEL on the MNS dataset, using four scales on top of the data scale. Compared to the ground truth labels, the overall best clustering was obtained for scale 3 with 24 cell clusters, having an ARI of 0.68, HS of 0.83 and CS of 0.84. Similar to the MNIST dataset, we checked the clustering result details by calculating the contingency matrix, showing indeed a good agreement (Supplementary Fig. S6), i.e. a clear one-to-one relation between SCHNEL clusters and ground truth labels. For example, cluster 10 with ‘Microglia’ and cluster 18 with ‘Olfactory ensheathing cells’. In some cases, SCHNEL merged similar classes into one cluster. For instance, cluster 1 contained two ‘Enteric’ cells (glia and neurons) classes. Cluster 23 grouped three classes of ‘Peripheral sensory neurons’, while cluster 15 had two classes of ‘Vascular cells’. Alternatively, SCHNEL did find two subtypes of ‘Astrocytes’ (cluster 3 and 7), and three subtypes of Oligodendrocytes (cluster 0,5 and 16).

## 4 Discussion

SCHNEL provides a scalable fast solution for clustering large single cell data. Its novel approach utilizes HSNE for informed sampling of the data points using the concept of landmark selection and area of influence, which preserves the manifold structure of the full data. The sampling reduces the computational challenge of clustering many data points to a problem of clustering a subset of the data points that is at least an order of magnitude smaller. The smaller subset can be quickly clustered by the Louvain algorithm, a graph-based clustering method. The manifold learning ensures that the sampled data points (landmark points) retain the same structure as the full dataset. As landmark points represent the data points, cluster labels can be easily propagated down the hierarchy to the full dataset.

Due to the informed sampling procedure, SCHNEL scales to large datasets. The results of the Phenograph-Data, HMIS and Panorama datasets showed that it is not necessary to cluster on the full dataset or even the most detailed scale in order to capture all clusters, even rare ones. In other words, a meaningful clustering can be obtained from a sampling of the data that is two, or more, orders of magnitude smaller than the full dataset. This gives SCHNEL the opportunity to cluster dataset sizes of up to millions of cells within workable timeframes.

When clustering the largest cytometry dataset, Phenograph-Data, SCHNEL was able to pool all cells from all individuals together in one clustering. Compared to a clustering of each individual separately (as done in the original Phenograph study), SCHNEL achieved two major advantages. Firstly, cell cluster frequencies across individuals can be directly applied as all clusters emerged from one clustering. This in contrast to clustering per individual which requires matching the clusters obtained across individuals to be able to compare their frequencies. Secondly, pooling all cells together helps to emphasize small rare cell populations, making them easier to detect. When analyzing per individual, rare cell population might be divided across individuals, resulting in too few cells to be detected as a separate population.

Using three CyTOF datasets, AML, BMMC, and Panorama, SCHNEL achieved similar or better performance compared to the tested existing tools. Moreover, SCHNEL did not require any pre-existing knowledge on how many clusters the data should contain. We observed an under-clustering of the BMMC dataset using SCHNEL, this may be due to the fact that the BMMC dataset contains 11 (out of 24) small cell populations with less than 1,000 cells. These small populations might not have enough representative landmarks in subsequent scales of the hierarchy.

When clustering the AML dataset, SCHNEL produced some interesting cell clusters that might have biological relevance. Clusters 2 and 3 separated the CD4 T-cells into two groups with different expression of CD7. CD7-CD4 T-cells have been reported before and can result from either ageing or prolonged immune system activation (Reinhold and Abken, 1997). Clusters 4 and 6 were only distinct in their expression of CD20. Both clusters mainly contained mature B cells. This suggests that cluster 4 (CD20-) is a plasma B cell population, as CD20 is known to be highly expressed across all mature B-cells except plasma cells (Leandro, 2013). Finally, Cluster 14 contained a set of 62 CD4 T-cells and 43 CD8 T-cells. Normally these two proteins are mutually exclusive in mature T-cells. It could be possible that SCHNEL detected a rare subset of CD4+CD8+ T-cells. Therefore, formation of this cluster seems mostly driven by high expression of CD235ab, a red blood cell protein used to filter them out. It is suggested that these cells were not properly filtered out during manual gating.

Clustering the MNS scRNA-seq dataset showed that SCHNEL can handle large feature dimensions and produce meaningful clustering. These result shows that SCHNEL can be used as general clustering tool for single-cell data, not only cytometry data.

It is important to note that SCHNEL is a stochastic procedure, as the HSNE employs an approximated nearest neighbor search for generating the transition matrix on the data scale. This means that, although extremely similar, different hierarchies made from the same data with the same parameters will be slightly different. In addition, the Louvain clustering algorithm is also stochastic because it chooses random nodes as candidates for merging when trying to optimize for modularity. Different runs of the Louvain method on the same graph may produce slightly different results.

The current implementation of SCHNEL provides clustering for all scales. This provides different level of details in the clustering, however, it limits the automation of the algorithm to produce one clustering of the data. Further improvements can automatically determine the scale providing the best clustering, which may also reduce the computation time.

In conclusion, SCHNEL presents a reliable automated clustering tool for single-cell high-dimensional datasets. Using the HSNE, SCHNEL allows to perform graph clustering scalable to tens of millions of cells. Such clustering can be applied at different scales of the hierarchy, providing different level of detail.

## Funding

This project was supported by the European Commission of a H2020 MSCA award under proposal number [675743] (ISPIC), the European Union’s H2020 research and innovation programme under the MSCA grant agreement No 861190 (PAVE), the NWO Gravitation project: BRAINSCAPES: A Roadmap from Neurogenetics to Neurobiology (NWO: 024.004.012), and the NWO TTW project 3DOMICS (NWO: 17126).

## Conflict of Interest

none declared.

## References

Aghaeepour, N. et al. (2013) Critical assessment of automated flow cytometry data analysis techniques. Nat. Methods, 10, 228–238.

Bandura, D.R. et al. (2009) Mass cytometry: Technique for real time single cell multitarget immunoassay based on inductively coupled plasma time-of-flight mass spectrometry. Anal. Chem., 81, 6813–6822.

Becher, B. et al. (2014) High-dimensional analysis of the murine myeloid cell system. Nat. Immunol., 15, 1181–1189.

Blondel, V.D. et al. (2008) Fast unfolding of communities in large networks. J. Stat. Mech. Theory Exp., 2008.

Chen, H. et al. (2016) Cytofkit: A Bioconductor Package for an Integrated Mass Cytometry Data Analysis Pipeline. PLoS Comput. Biol., 12.

Chester, C. and Maecker, H.T. (2015) Algorithmic Tools for Mining High-Dimensional Cytometry Data. J. Immunol., 195, 773–779.

Van Gassen, S. et al. (2015) FlowSOM: Using self-organizing maps for visualization and interpretation of cytometry data. Cytom. Part A, 87, 636–645.

Hernandez-Martinez, J.-M. et al. (2018) Interplay between immune cells in lung cancer: beyond T lymphocytes. Transl. Lung Cancer Res., 7, S336–S340.

Höllt, T. et al. (2016) Cytosplore: Interactive Immune Cell Phenotyping for Large Single-Cell Datasets. In, Computer Graphics Forum (Proceedings of EuroVis 2016).

de Koning, C. et al. (2016) Immune Reconstitution after Allogeneic Hematopoietic Cell Transplantation in Children. Biol. Blood Marrow Transplant., 22, 195–206.

Leandro, M.J. (2013) B-cell subpopulations in humans and their differential susceptibility to depletion with anti-CD20 monoclonal antibodies. Arthritis Res. Ther., 15.

Levine, J.H. et al. (2015) Data-Driven Phenotypic Dissection of AML Reveals Progenitor-like Cells that Correlate with Prognosis. Cell, 162, 1–14.

Maaten, L. van der (2014) Accelerating t-SNE using tree-based algorithms. J. Mach. Learn. Res., 15, 3221–3245.

van der Maaten, L. and Hinton, G. (2008) Visualizing Data using t-SNE. J. Mach. Learn., 9, 2579–2605.

Mosmann, T.R. et al. (2014) SWIFT-scalable clustering for automated identification of rare cell populations in large, high-dimensional flow cytometry datasets, Part 2: Biological evaluation. Cytom. Part A, 85, 422–433.

Newman, M.E.J. (2006) Modularity and community structure in networks. Proc. Natl. Acad. Sci. U. S. A., 103, 8577–8582.

Pezzotti, N. et al. (2017) Approximated and User Steerable tSNE for Progressive Visual Analytics. IEEE Trans. Vis. Comput. Graph., 23, 1739–1752.

Pezzotti, N. et al. (2020) GPGPU Linear Complexity t-SNE Optimization. IEEE Trans. Vis. Comput. Graph., 26, 1172–1181.

Pezzotti, N. et al. (2016) Hierarchical Stochastic Neighbor Embedding. In, Computer Graphics Forum (Proceedings of EuroVis 2016).

Picot, J. et al. (2012) Flow cytometry: Retrospective, fundamentals and recent instrumentation. Cytotechnology, 64, 109–130.

Rand, W.M. (1971) Objective criteria for the evaluation of clustering methods. J. Am. Stat. Assoc., 66, 846–850.

Reinhold, U. and Abken, H. (1997) CD4+CD7-T cells: A separate subpopulation of memory T cells? J. Clin. Immunol., 17, 265–271.

Rosenberg, A. and Hirschberg, J. (2007) V-Measure: A conditional entropy-based external cluster evaluation measure. In, EMNLP-CoNLL 2007 - Proceedings of the 2007 Joint Conference on Empirical Methods in Natural Language Processing and Computational Natural Language Learning., pp. 410–420.

Rundberg Nilsson, A. et al. (2013) Frequency determination of rare populations by flow cytometry: A hematopoietic stem cell perspective. Cytom. Part A, 83, 721–727.

Samusik, N. et al. (2016) Automated mapping of phenotype space with single-cell data. Nat. Methods, 13, 493–496.

Shekhar, K. et al. (2014) Automatic Classification of Cellular Expression by Nonlinear Stochastic Embedding (ACCENSE). Proc. Natl. Acad. Sci. U. S. A., 111, 202–207.

Spitzer, M.H. and Nolan, G.P. (2016) Mass Cytometry: Single Cells, Many Features. Cell, 165, 780–791.

Stikvoort, A. et al. (2017) Combining flow and mass cytometry in the search for biomarkers in chronic graft-versus-host disease. Front. Immunol., 8.

Traag, V.A. et al. (2019) From Louvain to Leiden: guaranteeing well-connected communities. Sci. Rep., 9.

van Unen, V. et al. (2016) Mass Cytometry of the Human Mucosal Immune System Identifies Tissue- and Disease-Associated Immune Subsets. Immunity, 44, 1227–1239.

Van Unen, V. et al. (2017) Visual analysis of mass cytometry data by hierarchical stochastic neighbour embedding reveals rare cell types. Nat. Commun., 8, 1–10.

Virgo, P.F. and Gibbs, G.J. (2012) Flow cytometry in clinical pathology. Ann. Clin. Biochem., 49, 17–28.

Weber, L.M. and Robinson, M.D. (2016) Comparison of Clustering Methods for High-Dimensional Single-Cell Flow and Mass Cytometry Data. Cytom. A, 89, 1084-1–96.

Zeisel, A. et al. (2018) Molecular Architecture of the Mouse Nervous System. Cell, 174, 999–1014.e22.

