## Supplementary Figures for "SCHNEL: Scalable clustering of high dimensional single-cell data"

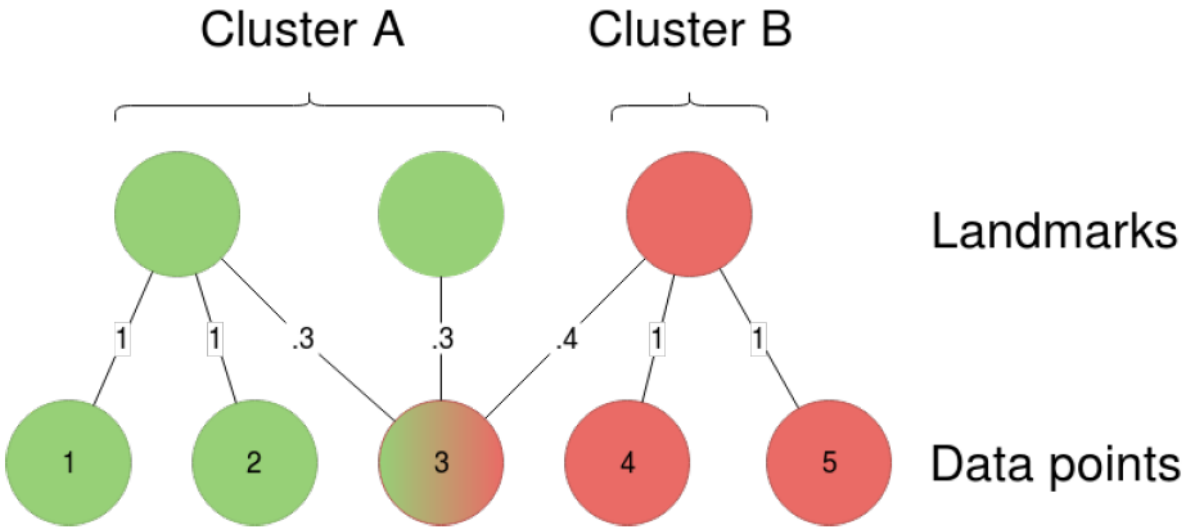


**Supplementary Fig. S1 Visual representation of propagating labels.** Three landmarks are clustered into two groups. The area of influence is represented by the lines between the landmarks and the data points. The numbers on the edges represent the probability of a data point being well presented by a landmark. Propagating the landmark labels down to the data level is unambiguous for data point 1, 2, 4, and 5. Point 3 will be assigned to cluster A since it has a higher total probability (0.3 + 0.3 = 0.6 versus 0.4)

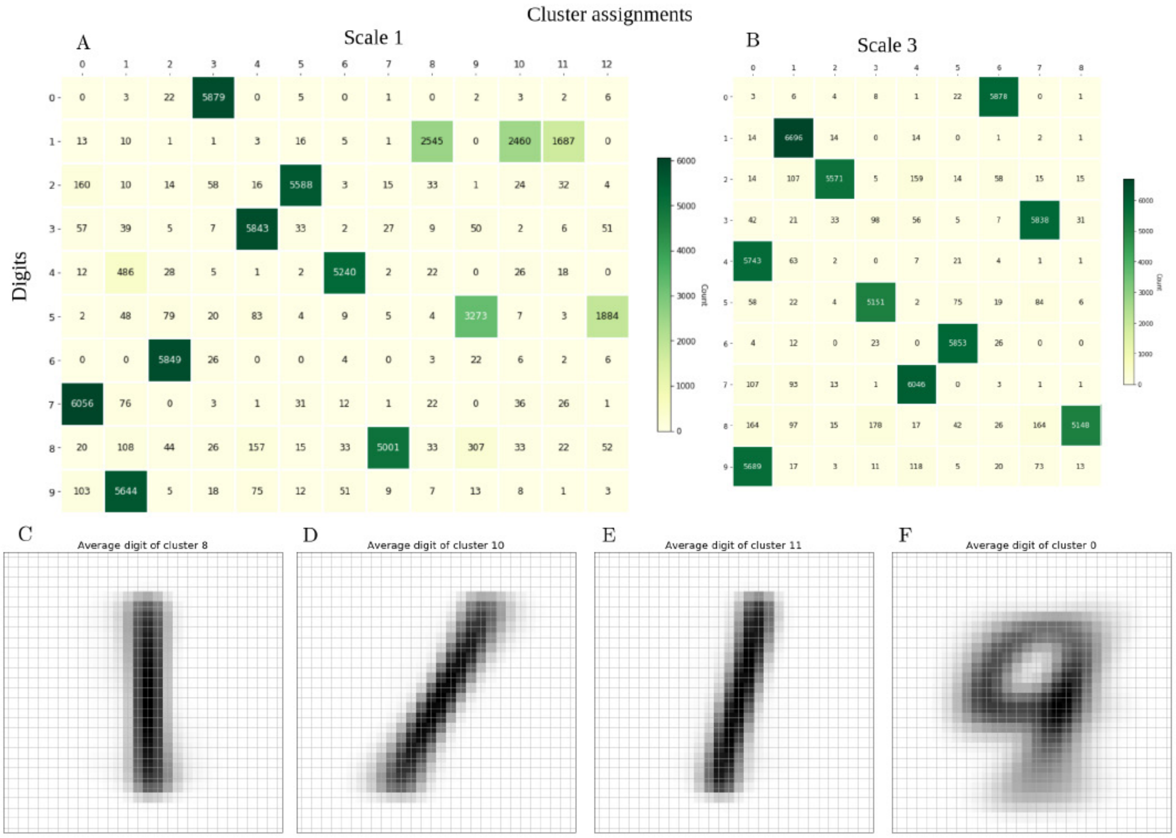


**Supplementary Fig. S2 (A)** Contingency matrix of clustering scale 1 of the MNIST dataset. Ones were separated into three different clusters, and fives were split over two different clusters. **(B)** Contingency matrix of clustering scale 3 of the MNIST dataset. All digits were assigned their own cluster, except fours and nines which were merged. **(C-E)** The average pixel values for the clusters containing ones in scale 1, clusters 8, 10, and 11, respectively. **(F)** The average pixel values for cluster 0 on scale 3, which contained fours and nines.

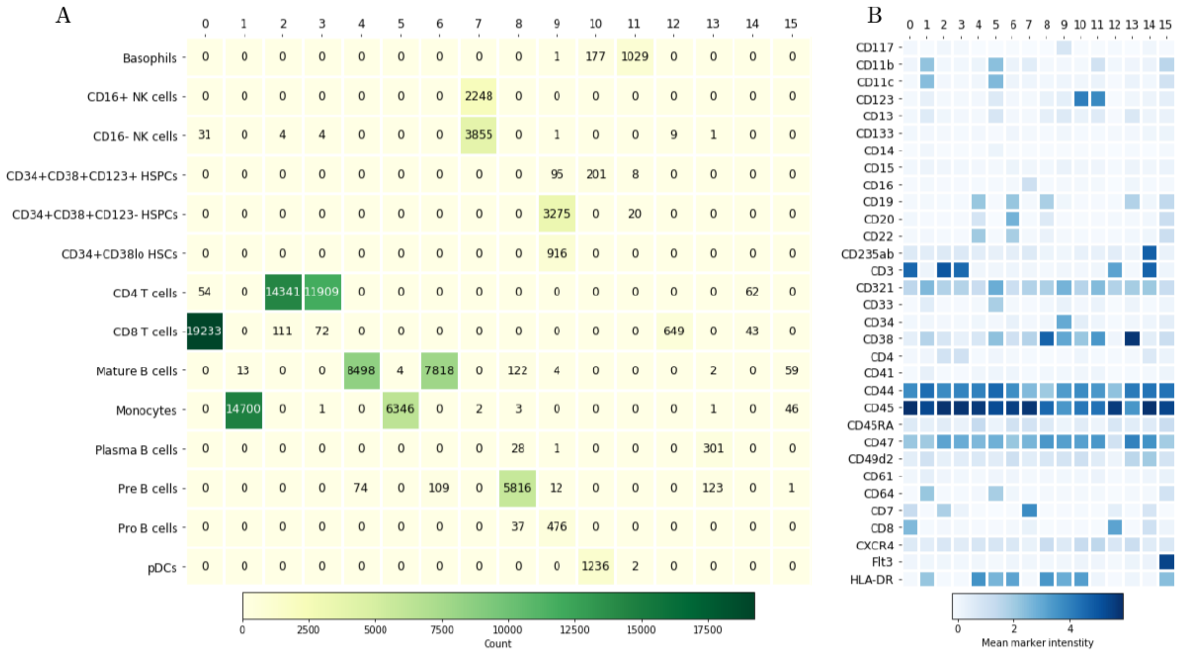


**Supplementary Fig. S3 (A)** Contingency matrix of scale 1 compared to manual annotations of the AML dataset. **(B)** Heatmap showing the average marker expression per cluster for the results of AML scale 1. Darker color means higher expression of that protein for that cluster.

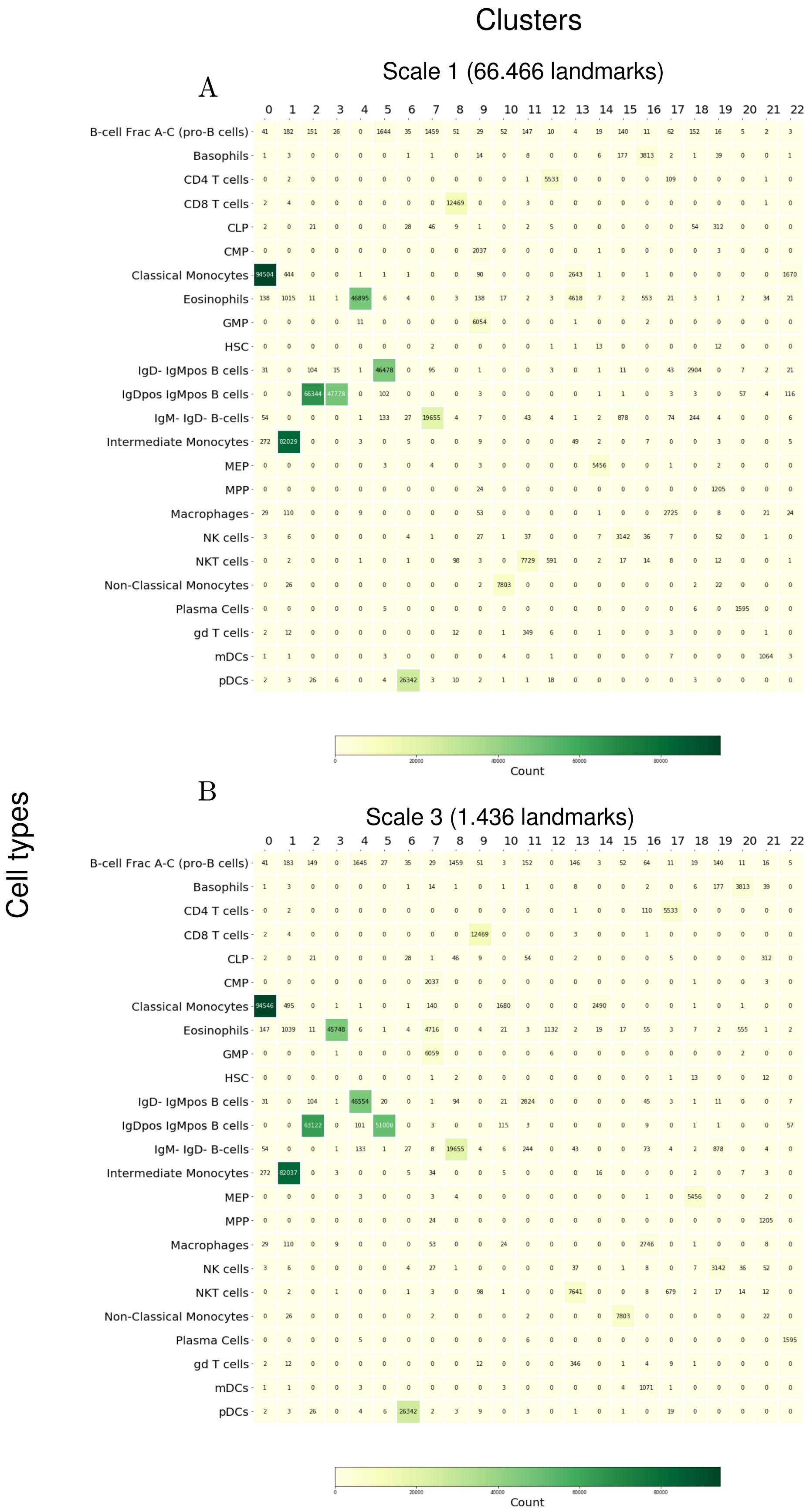


**Supplementary Fig. S4** Contingency matrices of the Panorama dataset clustering using SCHNEL on **(A)** scale 1 and **(B)** scale 3. Despite the very large difference in number of landmarks, the results are extremely similar.

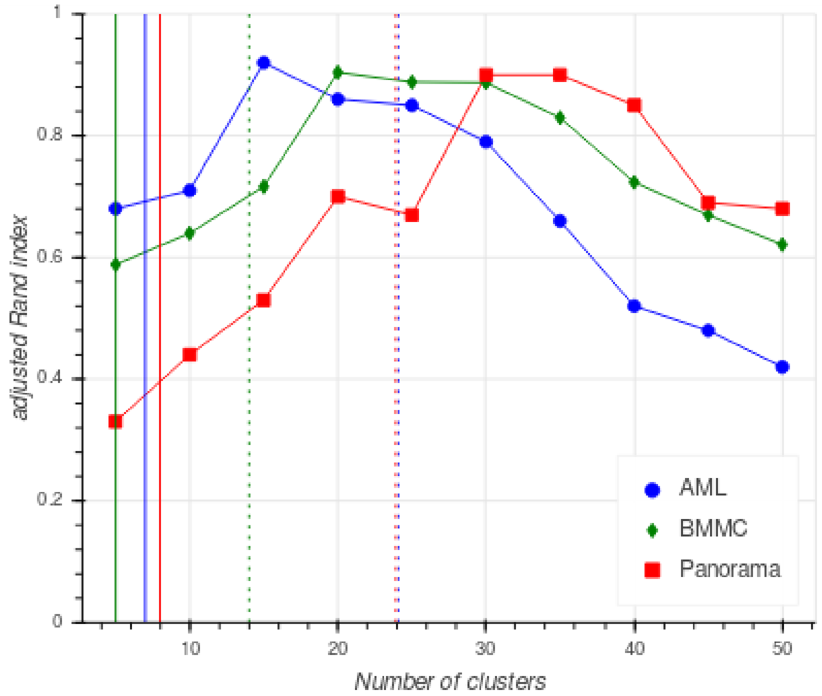


**Supplementary Fig. S5** Adjusted Rand index of FlowSOM on AML, BMMC, and Panorama datasets. Ten runs were performed for each dataset with a different forced number of clusters, ranging from 5 to 50 with increments of 5. The solid vertical lines are the number of clusters FlowSOM outputs when it was allowed to optimize the number of clusters automatically. The dashed vertical lines indicate the number of cell populations per the manual annotations.

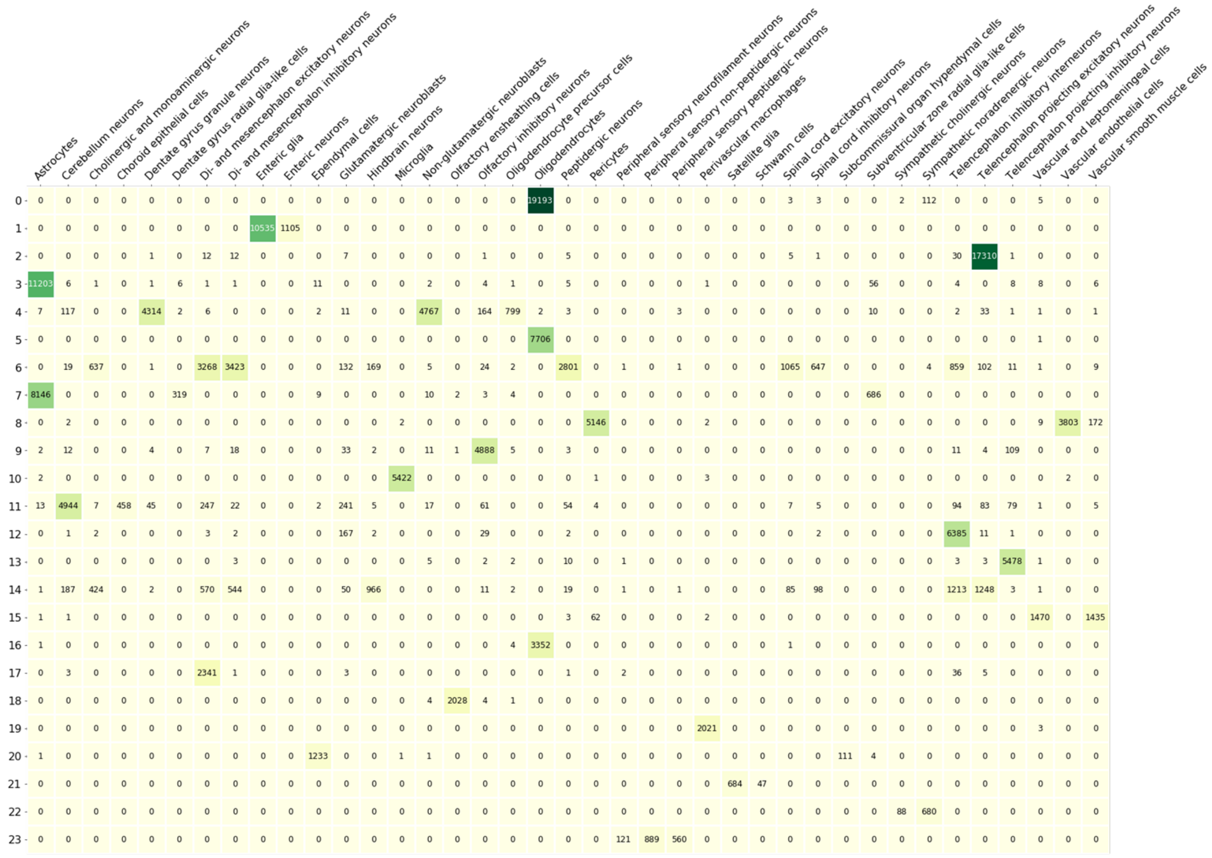


**Supplementary Fig. S6** Contingency matrix of the MNS dataset clustering using SCHNEL on scale 3.
